# Maternal dietary deficiencies in folates or choline during pregnancy and lactation worsen stroke outcome in 3-month-old male and female mouse offspring

**DOI:** 10.1101/2022.09.28.509960

**Authors:** McCoy Clementson, Lauren Hurley, Sarah Coonrod, Calli Bennett, Purvaja Marella, Agnes S. Pascual, Kasey Pull, Brandi Wasek, Teodoro Bottiglieri, Olga Malysheva, Marie A. Ca udill, Nafisa M. Jadavji

## Abstract

Maternal one-carbon (1C) metabolism plays an important role in early life programming. There is a well-established connection between the fetal environment and the health status of the offspring. However, there is a knowledge gap on how maternal nutrition impacts stroke outcome in offspring. The aim of our study was to investigate the role of maternal dietary deficiencies in folic acid or choline on stroke outcome in 3-month-old offspring. Adult female mice were fed a folic acid deficient diet (FADD), choline deficient diet (ChDD), or control diet (CD) prior to pregnancy. They were continued on diets during pregnancy and lactation. Male and female offspring were weaned onto a CD and at 2 months of age were subject to ischemic stroke within the sensorimotor cortex via photothrombosis damage. At 3-months-of-age, motor function was measured in offspring and tissue was collected for analysis. Mothers maintained on either a FADD or ChDD had reduced levels of *S*-adenosylmethionine in liver and *S*-adenosylhomocysteine in plasma. After ischemic stroke, motor function was impaired in 3-month-old offspring from deficient mothers compared to CD animals. In brain tissue, there was no difference in ischemic damage volume. When protein levels were assessed in brain tissue, there were lower levels of neurodegeneration in males compared to females and betaine levels were reduced in offspring from ChDD mothers. Our results demonstrate that a deficient maternal diet during critical timepoints in neurodevelopment results in worse stroke outcomes. This study emphasizes the importance of maternal diet and the impact it can have on offspring health.

## Introduction

Maternal deficiencies in one-carbon (1C) metabolism are known to have an impact on the neurodevelopment of offspring beginning with the closure of the neural tube during *in utero* development (1). Pregnant mothers who are deficient in folate have a higher incidence of birthing offspring with neural tube defects (NTDs) due to the high demand for nucleotide synthesis required by the early gestational period (2). Women of childbearing age are recommended to supplement with folic acid prior to conceiving, since the neural tube closes between 21 and 28 days after conception and most mothers do not know they are pregnant, on average, until 5.5 weeks (3). In response to this, the U.S. Public Health Service introduced mandatory folate fortification of grains in 1998, and the prevalence of NTDs, such as anencephaly and spina bifida, decreased by 28% (4, 5). Furthermore, pregnant, and lactating women are especially at risk for deficiencies in due to the physiological demands of the rapidly growing offspring. This is evidenced by the elevated requirements for an adequate intake (AI) of folic acid in those populations, which is 450mg/day for pregnant women and 550mg/day for lactating women, as compared to 425mg/day for non-pregnant women (2).

Folate, also known as vitamin B9, is the main component of 1C metabolism. Folates act as 1C donors in the regeneration of methionine from homocysteine, nucleotide synthesis and repair, and DNA methylation. Choline is another component of 1C metabolism and is critical for brain development. Studies have shown that during pregnancy and lactation choline stores get depleted (6, 7). Experiments using murine models have found that choline deficiencies during gestation result in numerous and concerning health consequences in the offspring. These experiments have shown that a choline deficiency *in utero* can cause abnormal sensory inhibition (8), affect spatial memory and hippocampal plasticity by altering DNA methylation patterns (9, 10) and induce hippocampal degeneration in the offspring. Thus, choline availability *in utero* has a part to play when it comes to cardiovascular health events. Yet, the severity of a choline deficient diet regarding stroke outcome is yet to be established and is an aim of our study.

There is a well-established connection between the maternal environment and the health of the offspring throughout the entirety of its life (11). Illustrating this point, one study demonstrates birthweights of half-siblings related by a common mother were correlated by an r-value of 0.58, compared to half-siblings related by a common father, whose correlation was 0.1 (12). Both low and high birth weights are associated with significant long-term health complications, including metabolic and cognitive development disorders (13–15). In addition, the health status of the mother has direct consequences on the offspring even once the offspring reaches adulthood and experiences a health crisis. As demonstrated in mice, offspring of mothers with gestational hypertension due to disrupted atrial natriuretic peptide (ANP) gene had worse outcomes following a stroke. When the offspring were subjected to a stroke of the middle cerebral artery, they had larger infarct volumes compared to the control (16). Nutrition is an important factor regulating the uterine environment, and deficiencies in the maternal diet will inevitably lead to deficiencies in the type of nutrients delivered to the placenta. For example, in pregnant baboons that were fed low-protein diets, the placenta downregulated amino acid transporters resulting in lower than typical levels of leucine and isoleucine (10).

Folate is required for the recycling of homocysteine to methionine. Without this conversion, the folate becomes trapped, which leads to hyperhomocysteinemia, a marker for the development of inflammation and cardiovascular disease (17). Cardiovascular disease is a leading cause of death worldwide, with stroke being the second most common cause (16). Stroke outcomes are varied between the ages of the victims, where the infarct occurs, the size of the infarct, and the degree of neurodegeneration that occurs post-infarct. At this time, it is unknown how folate and choline deficiencies *in utero* can affect the severity of a stroke event in the offspring. As it has been shown that maternal nutritional programming affects the health of the offspring throughout the course of its life, we aim to investigate how maternal dietary deficiencies in folic acid or choline *in utero* and during lactation will impact stroke outcome of 3-month-old male and female offspring.

## Materials and Methods

### Animals and experimental design

All experiments were conducted in accordance with the guidelines of the Midwestern University Institutional Animal Care Users Committee (IACUC 2983). Female and male C57/BL6J mice were obtained from Jackson laboratories. Male (n = 41) and female (n = 38) offspring were generated from breeding pairs.

Experimental manipulations are summarized in Figure 1. Two-month-old female mice were habituated for seven days before they were placed on either control (CD), folic acid (FD), or choline deficient diets (ChDD) (Table 1). The dams were maintained on the diets 4 weeks prior to pregnancy, during pregnancy, and lactation. Once the offspring were weaned from their mothers, they were maintained on a CD. At 2 months of age, the offspring were subjected to ischemic stroke, using the photothrombosis model to the sensorimotor cortex. Four weeks after damage, motor function was measured. After the completion of behavioral testing, brain tissue was collected for further analysis. Female breeding mice (dams) were euthanized after completion of pregnancies, plasma and liver tissue were collected for analysis of one-carbon metabolites.

**Figure 1.** Displays the timeline of the experiment. Pregnant mothers were fed either the CD, FD, or ChDD diet throughout the months of pregnancy and lactation until the offspring were weaned. Once the offspring were weaned, they were maintained on the CD. At 2 months of age, the offspring were subjected to ischemic stroke via the PT model. At 3 months, behavioral analysis was performed. At 3.5 months, animals were euthanized, and tissues were collected for various analyses.

**Table 1.**
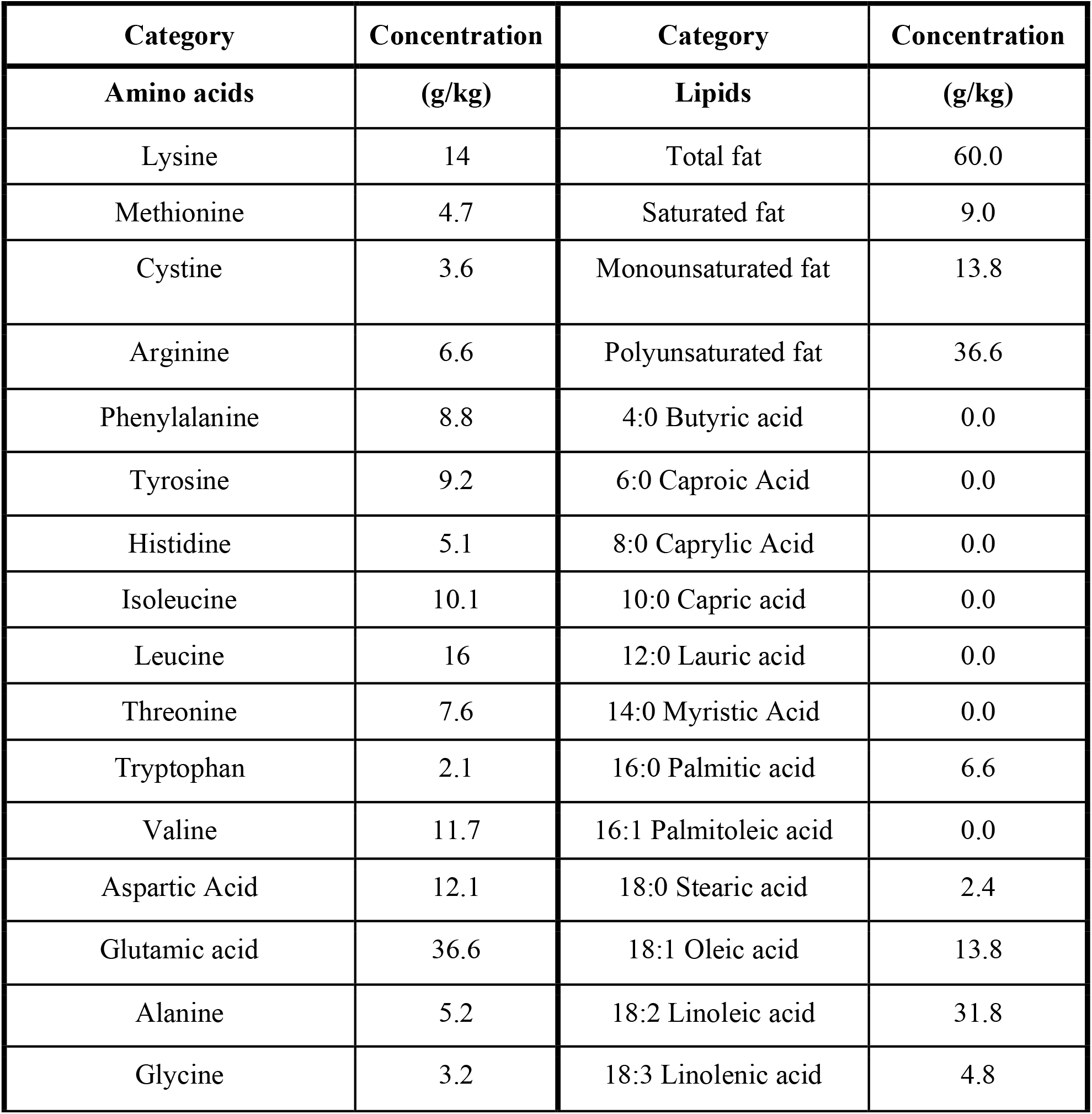
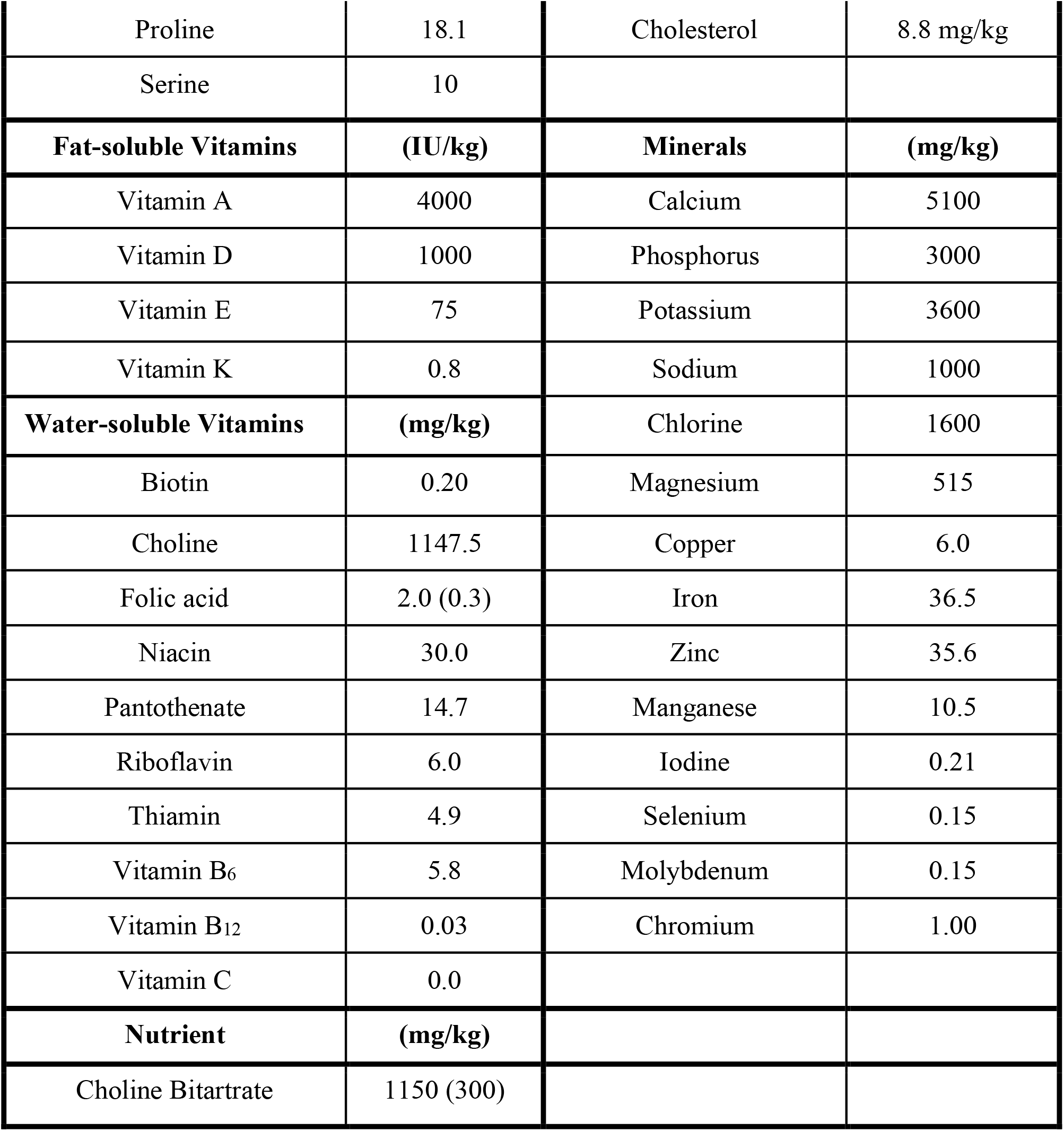
List of micro- and macro-nutrient contents in Envigo control (TD.01369) folic acid (TD.01546), choline (TD.06119) deficient diets. The Envigo folic acid deficient (TD.95247) was identical to the control diet (TD.04194) except it contained 0.2 mg/kg folic acid. The Envigo choline deficient (TD.06119) was identical to the control diet (TD.01369) except it contained 0.3 mg/kg choline bitartrate.

### Photothrombosis

At 2 months of age all female and male mice were subjected to photothrombosis to induce a unilateral ischemic stroke in the sensorimotor cortex. They were anesthetized with isoflurane (1.5%) in a 70:30 nitrous oxide: oxygen mixture. Core body temperature was monitored with a rectal thermometer (Harvard Apparatus) and maintained at 37 ± 0.2 ºC using a heating blanket. 10mg/kg of the photosensitive Rose Bengal dye was injected intraperitoneally 5 minutes prior to irradiation. A 532 nm green laser was placed 3 cm above the animal and directed to the sensorimotor cortex (mediolateral + 0.24mm) (18–20) for 15 min.

### Behavioral Testing

#### Accelerating Rotarod

An accelerating rotarod was used to assess balance and coordination. Mice were placed on a standard accelerating rotarod apparatus (Panlab Harvard Apparatus) which is 30cm above the ground, is 3 cm in diameter and 6 cm wide. The rotarod increases speed from 4 to 60 revolutions per minute over 8 minutes. When the mice fall, a digital sensor measures the latency to fall. Each mouse was subjected to three trials and the average was recorded (18).

#### Ladder Beam

To assess skilled motor function, mice were tasked to walk across a horizontal ladder. The ladder beam apparatus is two plexiglass walls with rungs in between spaced at irregular intervals, which are placed on two empty mouse cages. The time it takes the mouse to traverse the ladder is recorded. The animal is video recorded from below at a ventral angle so that all four paws can be visualized. Each step was scored on a scale of 0-6 by a blinded analyst. Scores of 0-2 are errors and the percent error was recorded as well as the overall movement score. Each mouse was subjected to 3 trials and the average of the percent error and movement score was used in statistical analysis (21).

#### Forepaw Placement

To further assess motor function, forepaw asymmetry was assessed. One mouse was placed in a vertical glass cylinder (19cm high, 14cm diameter) and the video was recorded above for 10 minutes. The first 20 movements were scored, and each contact of the forepaw against the cylinder was counted. The final score was calculated using the following formula: final score = (number of non-impaired forelimb movement − number of impaired forelimb movement)/(number of non-impaired forelimb movement + number of impaired forelimb movement + number of movements). A positive score indicates that the mouse prefers their non-impaired forelimb and a negative score indicate that the impaired forelimb is favored. Scores of 0 reflect equal use of each forelimb (22).

### Brain tissue

Brain tissue was sectioned using a cryostat (Fisher) at a thickness of 30μm and sections were slide mounted on microscope slides in serial order. One series of brain tissue was stained with cresyl violet (Sigma). Lesion volume was quantified using ImageJ (NIH) software (18, 23, 24).The lesion area was traced, and the area was multiplied by the section thickness and number of sections.

### Immunofluorescence

Brain tissue was used in immunofluorescence analysis to assess molecular mechanisms. Primary antibodies included, active caspase-3 (1:100, Cell Signaling Technologies) to measure apoptosis, hypoxia-inducible factor 1 alpha (HIF-1α, 1:100, AbCam), or choline acetyltransferase (ChAT, 1:100, Millipore). All brain sections were stained with a marker for neuronal nuclei, (NeuN, 1:200, AbCam). Primary antibodies were diluted in 0.5% Triton X and incubated with brain tissue overnight at 4°C. The next day, brain sections were incubated in Alexa Fluor 488 or 555 (Cell Signaling Technologies) and secondary antibodies were then incubated at room temperature for 2 hours and stained with 4’, 6-diamidino-2-phenylindole (DAPI) (1:1000, Thermo Fisher Scientific). The staining was visualized using a microscope (Zeiss) and all images were collected at the magnification of 400X.

In brain tissue within the ischemic core region, co-localization of active caspase-3, HIF-1α, or ChAT with NeuN labelled neurons were counted and averaged per animal. Images were merged, and a positive cell was indicated by co-localization of the antibodies of interest located within a defined cell. Cells were distinguished from debris by identifying a clear cell shape and intact nuclei (indicated by DAPI and NeuN) under the microscope. All cell counts were conducted by two individuals blinded to treatment groups. Using ImageJ, the number of positive cells were counted in three brain sections per animal. For each section, three fields were analyzed. The number of positive cells were averaged for each animal.

### One carbon metabolite measurement

Liver and plasma from female breeding mice (dams) was measured for total homocysteine, *S*-adenosylmethionine (SAM), *S*-adenosylhomocysteine (SAH), methionine, cystathionine, betaine, and choline using liquid chromatography tandem mass spectrometry (LC-MS/MS) as previously described (25).

Ischemic and non-ischemic cortical brain tissue from offspring was measured for acetylcholine, betaine, choline, glycerophosphocholine, methionine, phosphocholine, phosphatidylcholine, and sphingomyelin levels using the LC-MS method as previously reported (26, 27).

### Statistical Analysis

All data were analyzed by two individuals that were blinded to experimental treatment groups. Data was analyzed using GraphPad Prism 9.1.2. Two-way ANOVA analysis was performed comparing the mean measurement of both sex and dietary group for behavioral testing, plasma tHcy measurements, lesion volume, immunofluorescence staining, and choline measurements. If no sex differences were observed, male and female data was combined and a one-way ANOVA was run on behavioral testing, plasma tHcy measurements, lesion volume, immunofluorescence staining, and choline measurements. Significant main effects of one-way ANOVAs were followed up with Tukey’s post-hoc test to adjust for multiple comparisons. Statistical tests were performed using a significance level of 0.05. Data is presented as average ± standard error of the mean (SEM).

## Results

### Maternal dietary deficiencies result in SAM and SAH changes in liver and plasma tissue

We measured 1C metabolites in liver and plasma in females that were maintained on FADD, ChDD, and CD diets. In the liver tissue mothers maintained on FADD during pregnancy and lactation had reduced levels of SAM (Table 2; F(_2,17_) = 5.40, p = 0.02), whereas in the plasma FADD mothers had increased levels of SAH (Table 3; F(_2, 17_) = 4.65, p = 0.03).

**Table 2.** Maternal one-carbon metabolite concentrations in liver tissue. ^1^

| Metabolites<br>nmol/g | CD | FADD | ChDD | p-value |
| --- | --- | --- | --- | --- |
| Homocysteine | 20.9 ± 2.85 | 20.7 ± 2.28 | 27.0 ± 2.02 | 0.12 |
| S-adenosylmethionine | 57.9 ± 2.86 | 35.9 ± 5.47 | 44.4 ± 4.80 | 0.02 * |
| S-adenosylhomocysteine | 37.1 ± 3.46 | 44.2 ± 5.08 | 38.8 ± 2.59 | 0.93 |
| Methionine | 247.4 ± 18.1 | 248 ± 20.9 | 235 ± 12.4 | 0.20 |
| Cystathionine | 13.0 ± 1.94 | 14.0 ± 1.73 | 12.0 ± 3.31 | 0.15 |
| Betaine | 453 ± 91.4 | 394 ± 110 | 325 ± 33.4 | 0.63 |
| Choline | 484.9 ± 32.8 | 438 ± 28.8 | 485 ± 32.8 | 0.73 |

<sup>1</sup>Values are means ± SEMs. Abbreviations: CD, control diet; FADD, folic acid deficient diet; ChDD, choline deficient diet; L, liver; P, plasma. \* $p < 0.05$ , one-way ANOVA, diet effect.

**Table 3.** Maternal one-carbon metabolite concentrations in plasma tissue.^1^

| Metabolites | CD | FADD | ChDD | p-value |
| --- | --- | --- | --- | --- |
| Homocysteine ($\mu\text{M}$) | $9.21 \pm 0.95$ | $7.47 \pm 1.09$ | $9.27 \pm 0.772$ | 0.35 |
| S-adenosylmethionine (nM) | $280 \pm 13.1$ | $268 \pm 35.8$ | $253 \pm 18.8$ | 0.28 |
| S-adenosylhomocysteine (nM) | $161 \pm 46.9$ | $78.0 \pm 15.7$ | $46.3 \pm 7.62$ | 0.03* |
| Methionine ($\mu\text{M}$) | $84.4 \pm 5.76$ | $79.8 \pm 3.19$ | $76.0 \pm 10.3$ | 0.37 |
| Cystathionine (nM) | $942 \pm 142$ | $858 \pm 53.5$ | $908 \pm 128$ | 0.14 |
| Betaine ($\mu\text{M}$) | $58.5 \pm 4.72$ | $56.2 \pm 4.03$ | $55.4 \pm 6.1$ | 0.91 |
| Choline ($\mu\text{M}$) | $23.4 \pm 0.53$ | $19.4 \pm 2.01$ | $21.1 \pm 1.67$ | 0.27 |

<sup>1</sup>Values are means ± SEMs. Abbreviations: CD, control diet; FD, folic acid deficient diet; ChDD, choline deficient diet; L, liver; P, plasma. \* $p < 0.05$ , one-way ANOVA, diet effect.

### Impaired motor function in offspring after ischemic stroke

Four weeks after ischemic stroke to the sensorimotor cortex we measured motor function in female and male offspring using the accelerating rotarod and forepaw placement tasks. On the rotarod tasks, male and female offspring from FADD and ChDD mothers had reduced balance and coordination compared to CD animals (Figure 2A, (F (_2,59_) = 2.8, p < 0 .001). There was no effect of sex (F (_1,59_) = 0.30, p < 0.57). Performance on the forepaw placement task showed that there was also less impaired (Figure 2B, F(_2,48_) = 109.8, p < 0.001) and non-impaired (Figure 2C, F (_2,40_) = 9.5, p = 0.008) usage by male and female offspring from FADD and ChDD mothers. There was no effect of sex (impaired: (F (_1,48_) = 2.3, p < 0.14 and non-impaired: (F (_1,48_) = 1.59, p < 0.20).

**Figure 2.**
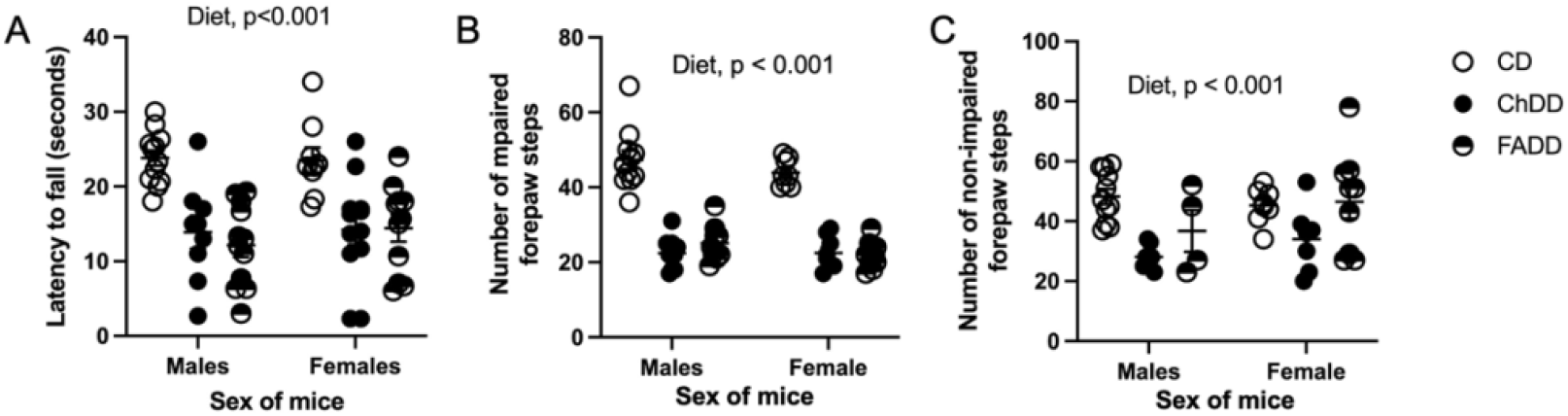
Motor function measurements in male and female offspring after ischemic stroke to the sensorimotor cortex. Using the accelerating rotarod we measured the latency to fall (A). The forepaw placement task measured usage of impaired (B) and non-impaired (C) forepaws. Depicted are means of ± SEM of 10 to 11 mice per group.

### No difference in ischemic damage volume in offspring

Five weeks after the induction of photothrombosis we measured ischemic damage volume in brain tissue of male and female offspring, representative images are shown in Figure 3A. Quantification of damage volume revealed that there was no difference in damage volume between maternal dietary groups (Figure 3B; F(_2,30_) = 0.80, p = 0.46) and sex (F(_1,30_) = 0.15, p = 0.70.

**Figure 3.** Impact of maternal dietary deficiency on ischemic stroke on damage volume. Representative cresyl violet image (A) and ischemic damage volume quantification (B). Depicted are means of ± SEM of 6 to 8 mice per group.

### Reduced neurodegeneration in male offspring

Within damaged ischemic tissue we assessed neurodegeneration in neurons by measuring active-caspase 3 levels. Representative images are shown in (Figure 4A) from offspring and semi-quantification was also conducted which revealed reduced degeneration in males compared to females (F(_1,11_) = 7.0, p = 0.02). There was no maternal diet effect (F(_2,12_) = 0.16, p = 0.85). We also measured hypoxia-inducible factor alpha (HIF-1α) within damage region and report a similar reduction neuronal cells of male offspring (Figure 4B; F(_1,14_) = 9.8, p = 0.007). There was no maternal diet effect (F(_2,14_) = 2.8, p = 0.10).

**Figure 4.** Impact of maternal dietary deficiencies on offspring neuronal apoptosis and response to hypoxia in an ischemic brain region. Representative active caspase-3, neuronal nuclei (NeuN) and 4′,6-diamidino-2-phenylindole (DAPI) staining. (A). Representative hypoxia-inducible factor alpha (HIF-1α), neuronal nuclei (NeuN) and DAPI staining. Quantification of HIF-1α, NeuN, and DAPI cell counts (B). Depicted are means of ± SEM of 3 mice per group. The scale bar = 50 μm. * p < 0.05, sex main effect.

### Offspring choline metabolites

We measured choline metabolites in cortical ischemic (Table 4) and non-ischemic (Table 5) brain tissue 4 weeks after the induction of ischemic stroke. Within ischemic cortical tissue we observed a decrease in betaine in FADD and ChDD offspring compared to CD animals (F(_2,31_) = 3.4, p = 0.04). Whereas in non-ischemic cortical tissue there was a sex difference (F(_1,31_) = 5.0 p = 0.03).

**Table 4.** Offspring one-carbon metabolite concentrations in ischemic (damaged) tissue. ^1^

| **Metabolites**<br>**nmol/g** | **CD** | **FADD** | **ChDD** | **p-values**<br>**Interaction**<br>**Sex**<br>**Diet** |
| --- | --- | --- | --- | --- |
| Methionine | Females<br>73.9 ± 4.3<br>Males<br>79.8 ± 3.7 | Females<br>71.6 ± 4.8<br>Males<br>69.1 ± 7.1 | Females<br>84.4 ± 3.8<br>Males<br>74.8 ± 3.1 | p = 0.28<br>p = 0.59<br>p = 0.14 |
| Acetylcholine | Females<br>3.9 ± 0.6<br>Males<br>4.0 ± 0.5 | Females:<br>3.8 ± 0.4<br>Males:<br>4.6 ± 0.7 | Females<br>3.7 ± 0.2<br>Males<br>3.7 ± 0.4 | p = 0.67<br>p = 0.46<br>p = 0.58 |
| Betaine | Females<br>15.3 ± 1.0<br>Males<br>16.0 ± 3.6 | Females<br>13.3 ± 0.97<br>Males<br>12.5 ± 1.9 | Females<br>10.6 ± 0.4<br>Males<br>11.6 ± 0.7 | p = 0.87<br>p = 0.86<br>p = 0.05* |
| Choline | Females<br>76.4 ± 7.9<br>Males<br>67.5 ± 4.3 | Females<br>67.9 ± 3.6<br>Males<br>67.5 ± 4.7 | Females<br>70.6 ± 5.2<br>Males<br>75.7 ± 6.4 | p = 0.45<br>p = 0.75<br>p = 0.57 |
| GPC | Females<br>529.8 ± 14.9<br>Males<br>540 ± 12.8 | Females<br>499.2 ± 35.3<br>Males<br>567.6 ± 33.3 | Females<br>517.4 ± 51.1<br>Males<br>530.4 ± 10.9 | p = 0.57<br>p = 0.23<br>p = 0.92 |
| Phosphocholine | Females<br>358.6 ± 15.3<br>Males<br>394.7 ± 10.5 | Females<br>346.1 ± 13.6<br>Males<br>386.3 ± 8.7 | Females<br>372.9 ± 19.5<br>Males<br>361.2 ± 16.9 | p = 0.16<br>p = 0.08<br>p = 0.73 |
| Phosphatidylcholine | Females<br>33421 ± 667.1 | Females<br>33171.3 ± 369.6 | Females<br>33171.3 ± 3.69.6 | p = 0.32<br>p = 0.44<br>p = 0.14 |
|  | Males<br>34804.6 ± 620 | Males<br>33394.8 ± 413.3 | Males<br>33394.8 ± 413.3 |  |
| Sphingomyelin | Females<br>4117 ± 182.1 | Females<br>4584.5 ± 144.4 | Females<br>4496.1 ± 244.0 | p = 0.32<br>p = 0.35<br>p = 0.07 |
|  | Males<br>4466.1 ± 139.5 | Males<br>4897 ± 279.4 | Males<br>4298.5 ± 171.1 |  |

<sup>1</sup>Values are means ± SEMs. Abbreviations: CD, control diet; FADD, folic acid deficient diet; ChDD, choline deficient diet; GPC, glycerophosphocholine.

**Table 5.** Offspring one-carbon metabolite concentrations in non-ischemic (healthy) brain tissue. ^1^

| Metabolites<br>nmol/g | CD | FD | ChDD | p-values<br>Interaction<br>Sex<br>Diet |
| --- | --- | --- | --- | --- |
| Methionine | Females<br>74.2 ± 5.8 | Females<br>73.8 ± 5.7 | Females<br>74.5 ± 6.8 | p = 0.77<br>p = 0.60<br>p = 0.73 |
|  | Males<br>80.2 ± 1.7 | Males<br>71.9 ± 7.4 | Males<br>77.7 ± 4.3 |  |
| Acetylcholine | Females<br>4.4 ± 0.6 | Females<br>4.9 ± 0.7 | Females<br>3.4 ± 0.3 | p = 0.28<br>p = 0.41<br>p = 0.40 |
|  | Males<br>4.1 ± 0.6 | Males<br>3.6 ± 0.4 | Males<br>3.9 ± 0.3 |  |
| Betaine | Females<br>14.4 ± 1.2 | Females<br>13.8 ± 0.5 | Females<br>12.4 ± 0.6 | p = 0.50<br>p = 0.11<br>p = 0.60 |
|  | Males<br>11.7 ± 1.3 | Males<br>12.7 ± 1.5 | Males<br>12.1 ± 0.9 |  |
| Choline | Females<br>86.1 ± 13.6 | Females<br>68.3 ± 4.4 | Females<br>69.4 ± 2.8 | p = 0.35<br>p = 0.70<br>p = 0.23 |
|  | Males<br>73.4 ± 4.5 | Males<br>67.4 ± 5.1 | Males<br>76.6 ± 5.4 |  |
| GPC | Females<br>572.5 ± 16.5<br>Males<br>553.4 ± 34.4 | Females<br>509.7 ± 32.6<br>Males<br>646.5 ± 24.0 | Females<br>493.7 ± 36.9<br>Males<br>557.7 ± 17.1 | p = 0.35<br>p = 0.25<br>p = 0.36 |
| Phosphocholine | Females<br>372.4 ± 15.3<br>Males<br>400.6 ± 20.0 | Females<br>355.2 ± 12.0<br>Males<br>376.9 ± 10.1 | Females<br>359.1 ± 14.2<br>Males<br>385.9 ± 10.7 | p = 0.97<br>p = 0.03*<br>p = 0.34 |
| Phosphatidylcholine | Females<br>34071.5 ± 624.8<br>Males<br>3478 ± 460 | Females<br>34698.8 ± 281.4<br>Males<br>34988.6 ± 713.3 | Females<br>33293.5 ± 167.4<br>Males<br>33780.1 ± 575.0 | p = 0.90<br>p = 0.23<br>p = 0.04 |
| Sphingomyelin | Females<br>4327.2 ± 215.5<br>Males<br>4692.7 ± 305.6 | Females<br>4704.8 ± 109.6<br>Males<br>4761.5 ± 132.6 | Females<br>4441.0 ± 182.3<br>Males<br>4702.3 ± 135.7 | p = 0.70<br>p = 0.15<br>p = 0.47 |

<sup>1</sup>Values are means ± SEMs. Abbreviations: CD, control diet; FAAD, folic acid deficient diet; ChDD, choline deficient diet; GPC, glycerophosphocholine. \* $p < 0.05$ , one-way ANOVA, main effect.

To further investigate the impact of maternal deficient diets and ischemic stroke on choline metabolism we measured levels of choline acetyltransferase within the ischemic brain region. Representative images are show in Figure 5 and semi-quantification revealed a sex difference (F(_1,12_) = 7.0, p = 0.02).

**Figure 5.**
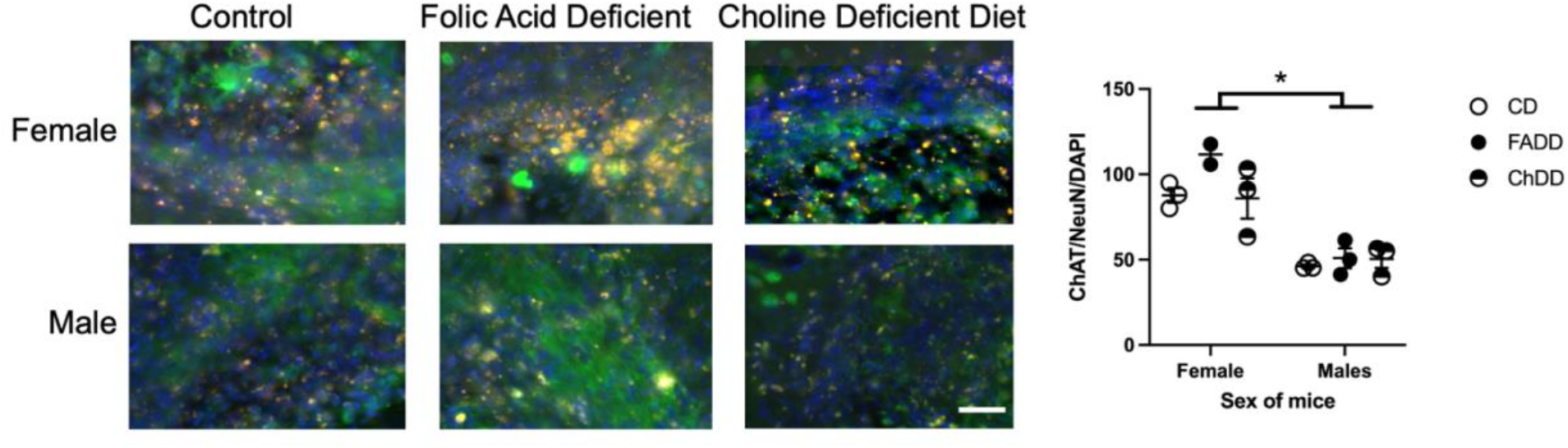
Impact of maternal dietary deficiency on ischemic stroke on choline acetyltransferase (ChAT) in ischemic damage brain region. Representative ChAT, neuronal nuclei (NeuN) and 4′,6-diamidino-2-phenylindole (DAPI) staining. Quantification of ChAT, NeuN, and DAPI cell counts (A). * p < 0.05, sex main effect.

## Discussion

Maternal nutrition during pregnancy and lactation impact offspring neurodevelopment, however the impact on offspring ischemic stroke outcome remains unclear. Our analysis of tissue metabolites revealed that decreased levels of SAMe in maternal liver indicating a lack of methylation capability and causing numerous downstream effects. Yet, in offspring only betaine and phosphatidylcholine are significantly reduced in offspring cortical ischemic and non-ischemic tissue, respectively, or simply put, most choline metabolites in cortical tissue were not significantly reduced in the experimental groups, FD or ChDD, as compared to the CD. In addition, the size of the ischemic damage volume between groups was not significantly different indicating that the nutrient deficiencies did not elicit a larger infarct size. Or rather, that the repair processes were not impacted by a lack of folic acid or choline. Furthermore, there were sex differences between the level of active caspase-3, hypoxia-inducible factor 1 alpha (HIF-1alpha), and choline acetyltransferase (ChAT) levels within ischemic brain tissue.

The health of the uterine environment is critical to the long-term health of the offspring (10–16, 28). While the fetus is often able to adapt to insults that occur *in utero*, however, these short-term adaptations can have life-long consequences (29). While folic acid is commonly known for its role in the closure of the neural tube, its effects on the vasculature are coming to light. Folic acid and its derivative 5-methyltetrahydrofolate have the ability to increase nitric oxide, leading to vasodilation, and scavenging free oxygen radicals, to decrease vascular injury (30). Given this, we posited that a deficiency in folic acid would have led to vasoconstriction and increased vascular damage post-ischemia. The analysis of the size and severity of the lesion volume suggests that this was not the case. However, while the internal measurements did not demonstrate any clear difference, the outward motor capabilities of the mice were significantly impaired as shown by the results of the rotarod task and the forepaw placement tasks. This could be explained by compensation of the nutrient-sufficient CD fed to the experimental groups post-weaning allowing the levels of metabolites to rise to normal levels but still hindering the development of the sensorimotor cortex to the point that motor function was still significantly impaired after the ischemic event. The degree to which micronutrient deficiencies *in utero* can affect the reparative ability of the brain post-stroke because of mistakes in early life programming when the demand for 1C metabolites is high.

Multiple studies have shown that females have higher levels of caspase and HIF-1alpha activation levels after ischemic stroke and our results are consistent with that (31, 32). However, given the higher degree of apoptosis indicated by those results, one could have expected to see sex differences in motor function as well. However, the female mice have higher levels of estrogen, which has a potent anti-inflammatory effect on the vasculature. For reference, post-menopausal women have much worse stroke outcome than premenopausal women (33). Studies in which female subjects have undergone ovariectomies have shown similar motor deficits compared to age-matched males (34). The decline in estrogen is a strong factor in determining the severity of outcome in female mice. Thus, it can be concluded that the presence of the neuroprotective estrogen in the young 3-month-old female mice could offset the higher degree of apoptosis and lead to similar motor function impairment.

Thus, it can be posited that folic acid supplementation should prevent certain adverse events from occurring by maintaining a mother’s blood folate within a specific healthy range. Dentists in Iran had the same idea and conducted a meta-analysis of research concerning the incidence of cleft lip and cleft palate correlated to the mother’s folic acid supplementation. They concluded that folic acid supplementation was significantly correlated with a decreased risk of both cleft lip and cleft palate^8^. In addition, experiments in murine models have revealed that folic acid supplementation can decrease adiposity, alleviate anxiety and depressive behavior, and improve neurogenesis^9–11^. In the case of ischemic stroke, our research group has shown that supplementation with 1C metabolites results in better stroke outcome (18, 19).

Whether or not choline supplementation can reverse any of these issues is an area of emerging research. A study in which mice were subjected to a folic acid deficient diet during gestation and a stroke event as an adult revealed the following results when their diet was supplemented with a B-vitamin cocktail of folic acid, choline, riboflavin, and choline after-birth: reduced sensorimotor deficits, decreased p53 expression, and increased plasticity of the perilesional area post-stroke event^18^.

Our results demonstrate that a deficient maternal diet in either folic acid or choline during critical timepoints in neurodevelopment (pregnancy and lactation) results in worse stroke outcomes. This study emphasizes the importance of maternal diet and the impact it can have on offspring health. Our study focused on 3-month-old offspring, it would be interesting to see if this continued into later adulthood as well as in old age.

## Abbreviations

(ChAT): Choline acetyltransferase
ChDD: Choline deficient diet
CD: control diet,
FADD: folic acid deficient diet
HIF-1alpha: hypoxia-inducible factor 1 alpha
SAM: *S*-adenosylmethionine
SAH: *S*-adenosylhomocysteine

## Acknowledgments

This research was funded by American Heart Association, grant number 20AIREA35050015 awarded to NMJ.

## References

1. Tulchinsky, T. H. (2010) Micronutrient Deficiency Conditions: Global Health Issues. Public Health Rev 32, 243–255

2. Osterhues, A., Ali, N. S., and Michels, K. B. (2013) The Role of Folic Acid Fortification in Neural Tube Defects: A Review. Critical Reviews in Food Science and Nutrition 53, 1180–1190

3. Ali, S. a and Economides, D. L. (2000) Folic acid supplementation. Current Opinion in Obstetrics and Gynecology 12, 507–512

4. Mills, J. L. and Signore, C. (2004) Neural tube defect rates before and after food fortification with folic acid. Birth Defects Research Part A: Clinical and Molecular Teratology 70, 844–845

5. Honein, M. A., Paulozzi, L. J., Mathews, T. J., Erickson, J. D., and Wong, L.-Y. C. (2001) Impact of Folic Acid Fortification of the US Food Supply on the Occurrence of Neural Tube Defects. JAMA 285, 2981–2986

6. Zeisel, S. (2004) Nutritional importance of choline for brain development. Journal of the American College of Nutrition 23, 621–626

7. Zeisel, S. H., Klatt, K. C., and Caudill, M. A. (2018) Choline. Advances in Nutrition 9, 58–60

8. Md, N., Cn, C., and Sh, Z. (2006) Dietary choline deficiency alters global and gene-specific DNA methylation in the developing hippocampus of mouse fetal brains. FASEB journal: official publication of the Federation of American Societies for Experimental Biology 20

9. I, P., Mj, K., Da, B., J, R., R, J., M, H., M, P., Jr, D., Aj, W., Pm, F., and De, J. (2014) Folic acid handling by the human gut: implications for food fortification and supplementation. The American journal of clinical nutrition 100

10. p, P., Fj, R., St, W., Pw, N., Tl, P., c, L., and t, J. (2016) Down-Regulation of Placental Transport of Amino Acids Precedes the Development of Intrauterine Growth Restriction in Maternal Nutrient Restricted Baboons. Biology of reproduction 95

11. Richmond, R. C., Sharp, G. C., Herbert, G., Atkinson, C., Taylor, C., Bhattacharya, S., Campbell, D., Hall, M., Kazmi, N., Gaunt, T., McArdle, W., Ring, S., Smith, G. D., Ness, A., and Relton, C. L. (2018) The long-term impact of folic acid in pregnancy on offspring DNA methylation: Follow-up of the Aberdeen folic acid supplementation trial (AFAST). International Journal of Epidemiology 47, 928–937

12. Mortensen, J. H. S., Øyen, N., Nilsen, R. M., Fomina, T., Tretli, S., and Bjørge, T. (2018) Paternal characteristics associated with maternal periconceptional use of folic acid supplementation. BMC Pregnancy and Childbirth 18, 1–8

13. Fan, R. G., Portuguez, M. W., and Nunes, M. L. (2013) Cognition, behavior and social competence of preterm low birth weight children at school age. Clinics (Sao Paulo) 68, 915–921

14. Nahum Sacks, K., Friger, M., Shoham-Vardi, I., Abokaf, H., Spiegel, E., Sergienko, R., Landau, D., and Sheiner, E. (2016) Prenatal exposure to gestational diabetes mellitus as an independent risk factor for long-term neuropsychiatric morbidity of the offspring. Am J Obstet Gynecol 215, 380.e1-7

15. Spiegel, E., Shoham-Vardi, I., Sergienko, R., Landau, D., and Sheiner, E. (2019) The association between birth weight at term and long-term endocrine morbidity of the offspring. J Matern Fetal Neonatal Med 32, 2657–2661

16. Ventura, N. M., Jin, A. Y., Tse, M. Y., Peterson, N. T., Andrew, R. D., Mewburn, J. D., and Pang, S. C. (2015) Maternal hypertension programs increased cerebral tissue damage following stroke in adult offspring. Molecular and Cellular Biochemistry 408, 223–233

17. Temel, S., Erdem, Ö., Voorham, T. A. J. J., Bonsel, G. J., Steegers, E. A. P., and Denktas, S. (2015) Knowledge on preconceptional folic acid supplementation and intention to seek for preconception care among men and women in an urban city: a population-based cross-sectional study. BMC Pregnancy Childbirth 15, 340

18. Jadavji, N. M., Emmerson, J., Willmore, W. G., MacFarlane, A. J., and Smith, P. (2017) B-vitamin and choline supplementation increases neuroplasticity and recovery after stroke. Neurobiology of Disease 103, 89–100

19. Jadavji, N. M., Mosnier, H., Kelly, E., Lawrence, K., Cruickshank, S., Stacey, S., McCall, A., Dhatt, S., Arning, E., Bottiglieri, T., and Smith, P. D. (2019) One-carbon metabolism supplementation improves outcome after stroke in aged male MTHFR-deficient mice. Neurobiology of Disease 132

20. Lee, J.-K., Kim, J.-E., Sivula, M., and Strittmatter, S. M. (2004) Nogo receptor antagonism promotes stroke recovery by enhancing axonal plasticity. The Journal of neuroscience: the official journal of the Society for Neuroscience 24, 6209–6217

21. Farr, T. D., Liu, L., Colwell, K. L., Whishaw, I. Q., and Metz, G. A. (2006) Bilateral alteration in stepping pattern after unilateral motor cortex injury: a new test strategy for analysis of skilled limb movements in neurological mouse models. Journal of neuroscience methods 153, 104–113

22. Theoret, J. K., Jadavji, N. M., Zhang, M., and Smith, P. D. (2016) Granulocyte macrophage colony-stimulating factor treatment results in recovery of motor function after white matter damage in mice. European Journal of Neuroscience 43

23. Jensen, E. C. (2013) Quantitative Analysis of Histological Staining and Fluorescence Using ImageJ. The Anatomical Record 296, 378–381

24. Jadavji, N., Emmerson, J. T., Shanmugalingam, U., Willmore, W. G., Macfarlane, A. J., and Smith, P. D. (2018) A genetic deficiency in folic acid metabolism impairs recovery after ischemic stroke. Experimental neurology 309, 14–22

25. Ducros, V., Belva-Besnet, H., Casetta, B., and Favier, A. (2006) A robust liquid chromatography tandem mass spectrometry method for total plasma homocysteine determination in clinical practice. Clinical Chemistry and Laboratory Medicine (CCLM) 44, 987–990

26. Holm, P., Ueland, P. M., Kvalheim, G., and Lien, E. A. (2003) Determination of Choline, Betaine, and Dimethylglycine in Plasma by a High-Throughput Method Based on Normal-Phase Chromatography – Tandem Mass Spectrometry. Clin Chem 49, 286–294

27. Taesuwan, S., McDougall, M. Q., Malysheva, O. V., Bender, E., Nevins, J. E. H., Devapatla, S., Vidavalur, R., Caudill, M. A., and Klatt, K. C. (2021) Choline metabolome response to prenatal choline supplementation across pregnancy: A randomized controlled trial. FASEB J 35, e22063

28. Virdi, S. and Jadavji, N. M. (2022) The Impact of Maternal Folates on Brain Development and Function after Birth. Metabolites 12, 876

29. Blackmore, H. L. and Ozanne, S. E. (2015) Programming of cardiovascular disease across the life-course. Journal of Molecular and Cellular Cardiology 83, 122–130

30. Stanhewicz, A. E. and Kenney, W. L. (2017) Role of folic acid in nitric oxide bioavailability and vascular endothelial function. Nutr Rev 75, 61–70

31. Zampino, M., Yuzhakova, M., Hansen, J., McKinney, R. D., Goldspink, P. H., Geenen, D. L., and Buttrick, P. M. (2006) Sex-related dimorphic response of HIF-1α expression in myocardial ischemia. American Journal of Physiology-Heart and Circulatory Physiology 291, H957–H964

32. Liu, F., Li, Z., Li, J., Siegel, C., Yuan, R., and McCullough, L. D. (2009) Sex Differences in Caspase Activation After Stroke. Stroke 40, 1842–1848

33. Koellhoffer, E. C. and McCullough, L. D. (2013) The Effects of Estrogen in Ischemic Stroke. Transl. Stroke Res. 4, 390–401

34. Liu, F., Yuan, R., Benashski, S. E., and McCullough, L. D. (2009) Changes in Experimental Stroke Outcome across the Life Span. J Cereb Blood Flow Metab 29, 792–802

